# shRNA drop-out screen identifies BRD4 targeting transcription from RNA polymerase II system to activate β-catenin to promote soft-tissue tumor proliferations

**DOI:** 10.1101/2023.07.14.548690

**Authors:** Sylvia Y. Sun, Vlad Tsiperson

**Affiliations:** Mortimer B. Zuckerman Research Center - Sloan Kettering Institute, 417 E 68th St, New York, NY 10065; Memorial Sloan Kettering Cancer Center, 1275 York Avenue, New York, NY 10065, USA

## Abstract

BRD4 (Bromodomain containing protein 4) is a chromatin reader binds to acetylated lysine residues on histones interacting with RNA Pol II, p-TFeb. PDGF-BB was presented here, in soft-tissue tumor, as an oncogenic factor driving cell proliferation, and aberrant BRD4 knockdown significantly reduced tumor aggressiveness and unfavorable prognosis in soft-tissue tumors. To identify suppressive key drivers impeding demoid tumor growth, shRNA drop-out screen analysis identified signature of “transcription from RNA polymerase II promoter” including DDX, Stat3, SMARCA, ATM, SIRT1, cMyc that were recruited with BRD4 interation in activating β-catenin, which is a major key driver mutated in soft-tissue tumor, and its depletion ceased soft-tissue tumor cell growth. Sepcifically, BRD4 mediated PDGF-BB signaling in GSK stimulation through transcriptional regulation from RNA polymerase II activity with PI3K as target, and thus not only canonical β -catenin/TCF4 signaling, but also non-canonical β -catenin conjunction complex response was activated by BRD4 in nucleus involved in promoting cell proliferation. Our study delineated a signaling axis that may allow soft-tissue tumor cells to escape apoptosis during colonization by activating PDGFBB-BRD4-GSK-β -catenin and non-canonical-β-catenin pathway through BRD4 in cancer cells. An efficient treatment for soft-tissue tumors could be accomplished by targeting PDGF and BRD4 survival pathways on soft-tissue tumor cells.

## Introduction

Soft-tissue tumors are a type of tumor that originate from fibrous tissue. Cells usually do not metastasize, however, local aggressive growth can cause significant morbidity by infiltrating surrounding structures with pain in patients’ limbs. Both sporadic and familial adenomatous polyposis (FAP) types of tumors are linked to constitutive activation of the Wnt signaling pathway with mutations in the β-catenin oncogene *CTNNB1* or the tumor suppressor gene *APC*, respectively. Radiotherapy is an effective modality for soft-tissue tumors, either alone or as an adjuvant to resection. Besides, other systemic options include more classic chemotherapeutic compounds that also play an effect.

It is now well established that BET proteins are frequently deregulated in cancer and contribute to aberrant chromatin remodeling and gene transcription that mediates tumorigenesis [1,2], even though the BET family of proteins were initially recognized for their role as important epigenetic regulators in inflammation and inflammatory diseases. The BET family consists of BRD2, BRD3, BRD4 and BRDT. They share two N-terminal bromodomains and an extra C-terminal domain, and their bromodomains show high levels of sequence conservation. Although BET bromodomains share a highly similarity of amino acid sequence, the transcriptional regulatory complexes recruited by each member are different. BRD4 is universally expressed in a variety of tissues[3,4], and it is capable of recruiting positive transcription elongation factor complexes (P-TEFb) and regulates DNA transcription of eukaryotes by RNA polymerase II (RNA Pol II)[5,6]. In addition, the ET domain of BRD4 can recruit transcription modifiers, such as histone methyltransferase NSD3 containing the SET domain, independently[7].

Aberrant expression of BRD4, promotes the progression of cell cycling, invasion and metastasis of cancer cells[8–10]. There is little direct study about the association between soft-tissue tumors and BRD4. Our lab recently performed a loss-of-function screening using shRNA pool, and there were some essential genes that are important for soft-tissue tumor cell growth — BRD4 was one of the genes. There are some non-direct evidence showed that targeting BRD4, or its specific downstream target, making BRD4 a potential therapy for human soft-tissue tumors[11,12].

## Results

### (#1). PDGF-BB promoted soft-tissue tumor proliferation associated with BRD4 and β-catenin

PDGF-BB promoted soft-tissue tumor proliferation (Figure 1a), and β-catenin was mutated at S45F[13]. β-catenin was regarded as a driver gene, and its knocked down directed the induction of cell death in primary FAP-associated tumor cells in a culture[14]. Chip-Seq data showed that PDGFR was significantly upregulated in *CTNNB1* knocked down cells, suggesting that β-catenin down-regulated PDGFR (data not shown).

**Figure 1.**
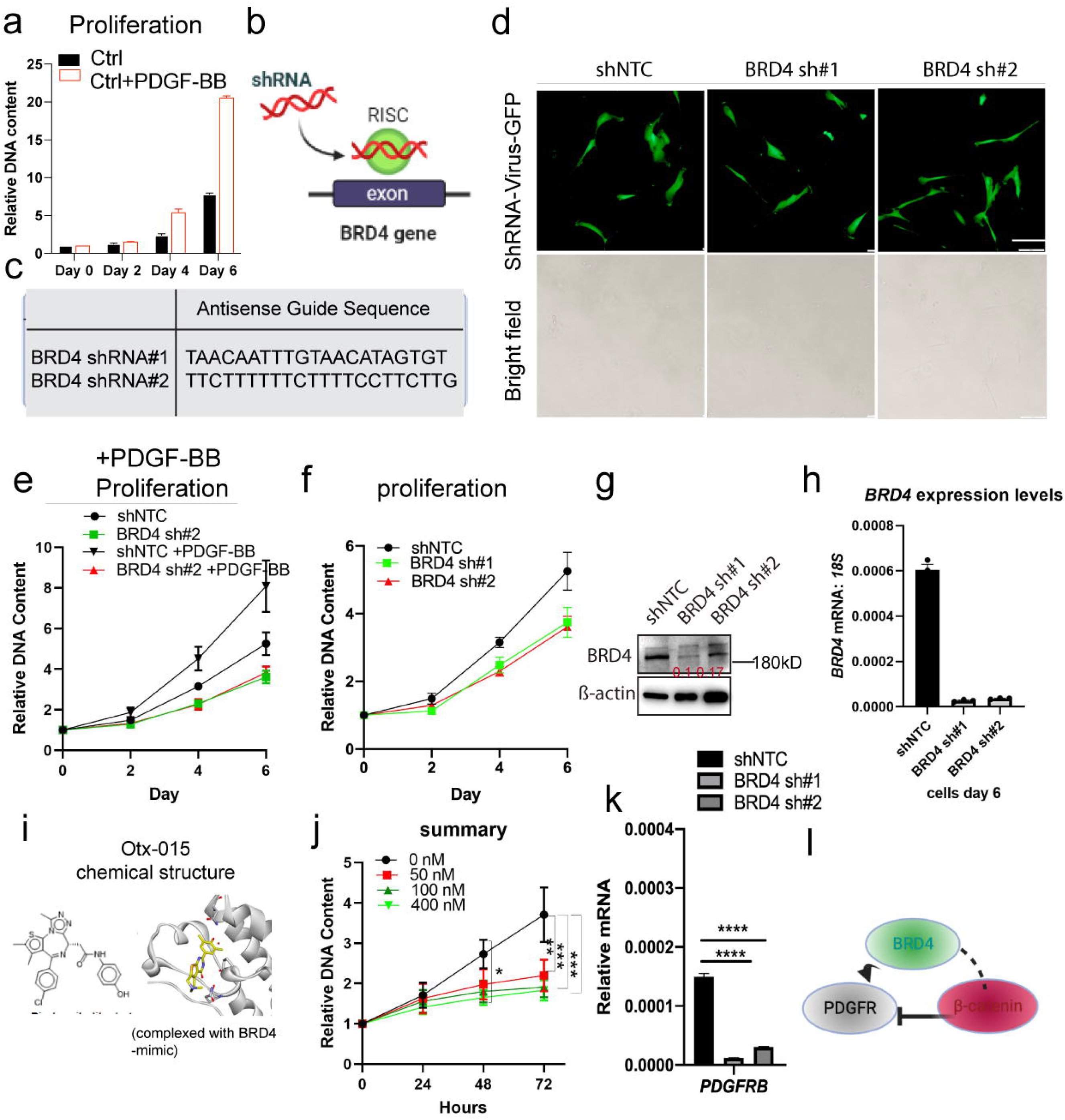
PDGF-BB promoted soft-tissue tumor cell proliferation associated with BRD4 and β-catenin. (a)proliferation assay performed over soft-tissue tumor cells with and without presense of PDGF-BB. (b).Schematic representation of the specific shRNA editing strategy. ShRNA was designed to specifically target the BRD4 allele. (c)The shRNA antisense guide sequence corresponds to the target BRD4 allele in human Ds soft-tissue tumor cells. (d)The shRNA virus infection efficiency. (bar, 10 μm) (e) Cell proliferation ability of BRD4 knock-down cells, and control cells without the presence of PDGF-BB, and in the presence of PDGF-BB. Mean ± SEM, n=4 independent result. (f).Cyquant cell proliferation assay in BRD4 knock-down cells (BRD4 sh#1, BRD4 sh#2) compared to control cells (shNTC). One-way ANOVA; ***p < 0.001,*p < 0.05, Mean ± SEM, n=4 independent result. (g).BRD4 expression in BRD4 knocked down cells by Western blot. (h).Relative mRNA levels of BRD4 by Real-time q-PCR analysis. (i)Inhibitor Otx-015 (Birabresib) chemical structure. (j) Cyquant cell proliferation assay in cells following inhibitors with different concentrations.One-way ANOVA; ***p < 0.001,*p < 0.05, Mean ± SEM, n=3 independent result. (k). PDGFR mRNA was regulated by BRD4. (o). An illustration to show that PDGF singaling was associated with BRD4, and β-catenin.

To describe the tumorigenic roles of BRD4 in soft-tissue tumors, two shRNA sequences targeting human BRD4 were designed and shRNA viruses with doxycycline (Dox)-induced tet-on system were delivered into human soft-tissue tumor Ds cells (Figure 1b, 1c). Cells were infected with shRNA virus with GFP induced(Figure 1d), suggesting a successful infection rate. It has been demonstrated that PDGFs participate in the proliferation, angiogenesis, migration, and invasion of many tumors, especially malignant tumors [15]. Our results showed that BRD4 knock-down induced weak proliferation of soft-tissue tumor, however, this phenotype could not be rescued by PDGF-BB (Figure 1e). This suggests that PDGF-BB induces soft-tissue cell proliferation with a requirement of BRD4. An impaired cell proliferation was observed in cells with BRD4 knockdown as measured by CyQuant assay (Figure 1f). BRD4 was knocked down by BRD4 shRNA-1 (BRD4 sh#1), and BRD4 shRNA-2 (BRD4 sh#2) compare to control (shNTC) at mRNA transcription level after doxycycline induction. Figure 1g showed that the protein abundance, and Figure 1h showed RNA of BRD4 were greatly reduced, thus verifying the specificity of knockdown. Following BRD4 inhibitor Otx-015 (Birabresib)’s exposure, which can be complexed with BRD4 using its BD domains (Figure 1i), impaired cell proliferation was evident, suggesting the potential synergic effects of Otx-015 (Figure 1j). Moreover, PDGFR’s mRNA level was significantly down-regulated in BRD4_kd Ds cells compared to control (Figure 1k). Overall, these data suggest that soft-tissue tumor cells with BRD4 inhibition reduces cell proliferation in soft-tissue tumors, and PDGF-BB in promoting soft-tissue tumor is associated with β-catenin and BRD4(Figure 1l).

### (#2). Inhibition of BRD4 resulted in cell cycle arrest, apoptosis accompanied by variance changes of morphology in soft-tissue tumor

To investigate the role of BRD4 knock-down on metastatic capability, we noticed that the cells with BRD4 knocked-down acquired a spindle-shaped morphology (Figure 2a). When cell length was measured, BRD4 knocked-down cells were more elongated compared to controls (Figure 2b). Cell cycle assay showed BRD4 knocked down resulted in cell cycle arrest in G0/G1(Figure 2c, 2d). pRb, as a marker of cell-cycle and apoptosis, was increased by BRD4 inhibition (Figure 2e). These data suggest that suppression of BRD4 induces cell cycle arrest accompanied by variance changes of morphology in soft-tissue tumor.

**Figure 2.**
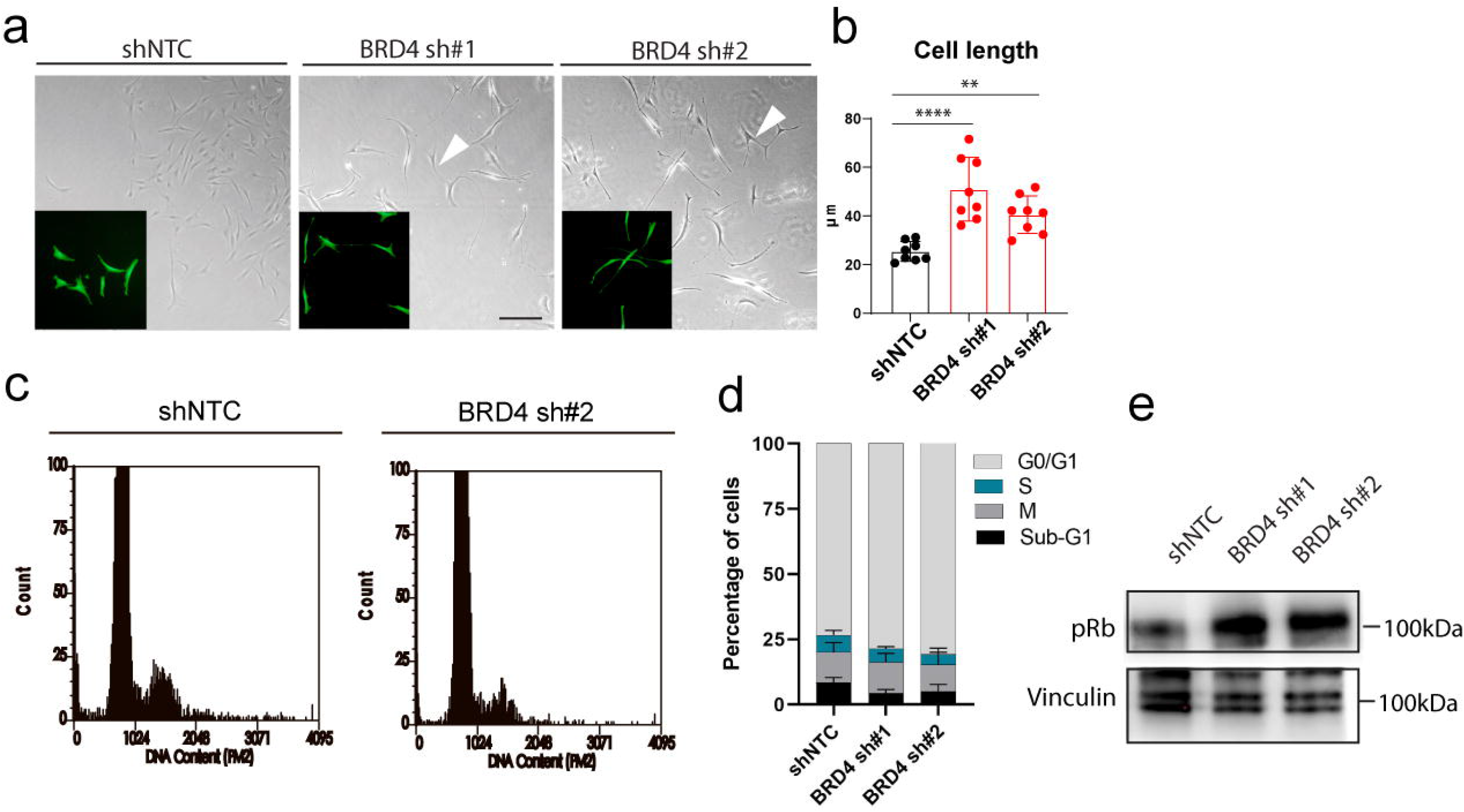
BRD4 inhibition causes morphology change, cell cycle arrest, and apoptosis of soft-tissue tumor cells, independently of p53. (a)In light microscopy, the morphology of soft-tissue tumor cells was studied. ShRNA editing of BRD4 knock-down induced differential changes. A spindle-shape was observed in the BRD4 knock-down cell shape. Scale bar, 10 μm.(b)The cell length of cells was calculated. Student’s t test; n=8. (c)Cell cycle assay based on BRD4 knock-down cells and control cells. (d)Analysis of cell cycle. One-way ANOVA; *p < 0.05, Mean ± SEM, n=5-6 independent samples. (e)Representative western showed increased pRb protein in BRD4 knock-down cells compared to control cells. Blot represent as least two biological repeats.

### (#3). BRD4 knock-down decreased active β-catenin and its targets

BRD4 knocked down showed a significant down-regulation of β-catenin (Figure 3a) at protein level (Figure 3b) and mRNA level (Figure 3c). Its nucleus downtream target HDAC2 was also down-regulated suggesting the active form of β-catenin was also regulated by BRD4 (Figure 3d). The complex partner TCF4 was downregulated under BRD4_kd condition, and the traditional Wnt/β-catenin target Wnt 7a was up-regulated since there was absense of Wnt (Figure 3e). These indicates that BRD4 regulates β-Wnt/catenin/TCF4 signaling. A known upstream activator of β-catenin FOXQ1 was down-regulated suggesting upstream β-catenin signaling was downregulated in BRD4 inhibition condition (Figure 3f).

**Figure 3.**
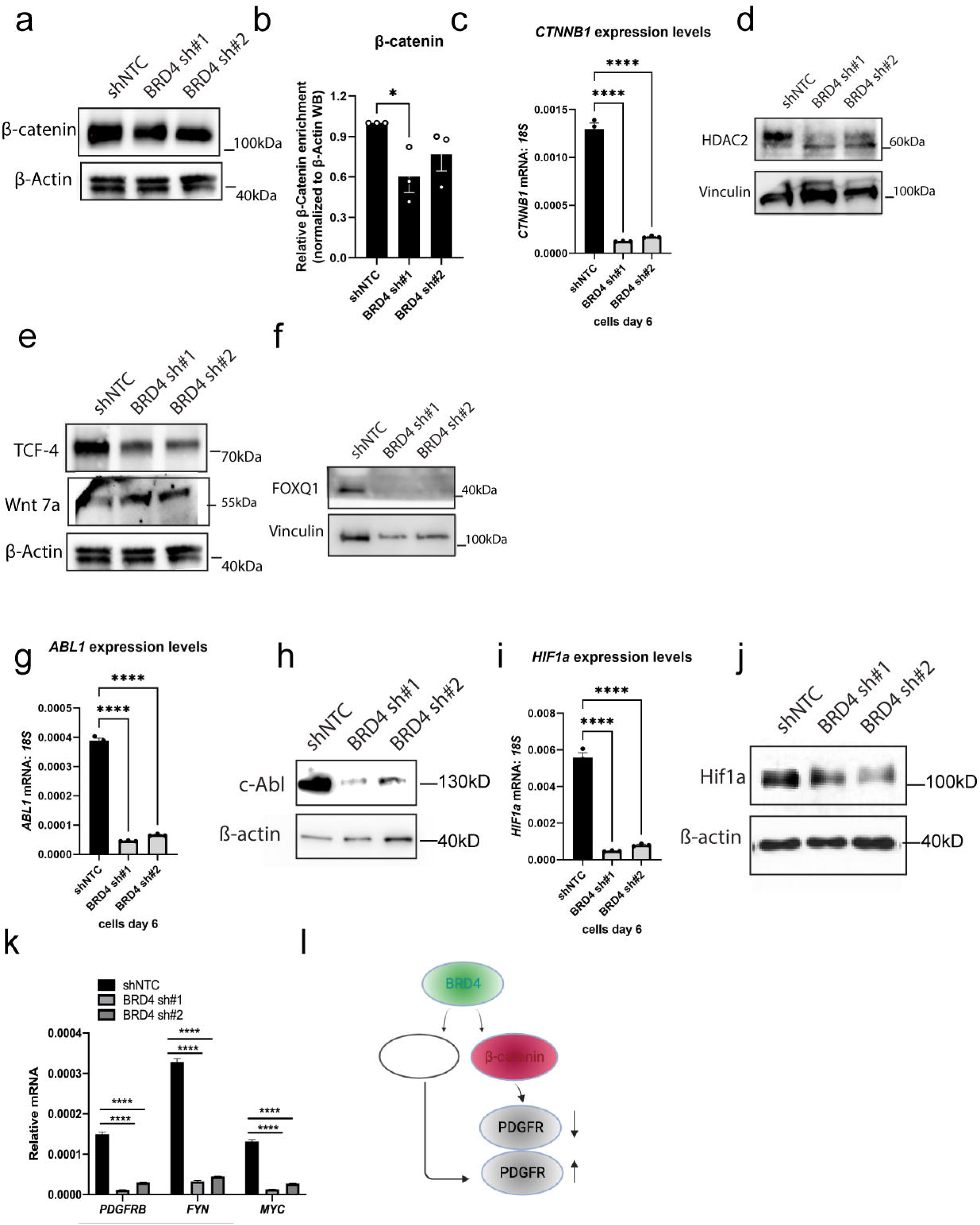
Knock-down of BRD4 induced significant reduction of β-catenin and its target, but BRD4 may also regulate PDGF signaling as upstream of β-catenin. .Down regulateion of beta-catenin in BRD4 knock-down cells compared to controls. (b).BRD4 induced 0.4 fold decrease of β-catenin protein compared to control. One-way ANOVA; *p < 0.05, Mean ± SEM, n=3 independent experiments. (c). Analysis of *CTNNB1* gene expression. One-way ANOVA; ****p < 0.0001, Mean ± SEM, n=3 independent samples. (d) HDAC2 expression under BRD4_kd condition. (e) β-catenin complex partner TCF4 expression and its target Wnt 7a. Blot represent as least two biological repeats. (f). FOXQ1 expression. Blot represent as least two biological repeats. (g,i)Analysis of gene expression. One-way ANOVA; ****p < 0.0001, Mean ± SEM, n=3 independent samples.(h,j). β-catenin target expression. (k)Analysis of other beta-catenin target gene expression. One-way ANOVA; ****p < 0.0001, Mean ± SEM, n=3 independent samples. (l) A diagram to show that BRD4 regulated β-catenin, but may also regulate PDGF signaling.

The inhibition of BRD4 strongly inhibited gene transcription of endogenous target, identified in β-catenin knocked down soft-tissue tumor Chip-Seq, at both transcription (Figure 3g, 3i) and protein level (Figure 3h, j), respectively. Consistently, other potential β-catenin target genes (identified in Chip-Seq, which may represent non-canonical β-catenin target/signaling): *PDGFR*, *FYN*, and *MYC* transcription were also significantly decreased (Figure 3k). However, gene pattern from Chip-Seq data from *CTNNB1* knocked down cells was different: at least *PDGFRB*, and *FYN* were opposingly regulated by BRD4 knock-down compared to *CTNNB1* knocked down, suggesting that BRD4 maintains β-catenin but PDGF signaling may be regulated by BRD4 as upstream signaling of β-catenin (data not shown). Together, these results indicate that BRD4, using alternative pathways, functions as a key regulator of mutated β-catenin for soft-tissue tumorigenesis.

### (#4). BRD4 inhibition disrupted PDGF-BB by decreasing phopho-GSK

β-catenin is degraded by a destruction complex which contains β-catenin, GSK, CK1[16], and GSK-3β works as upstream effector for β-catenin, MDM2, and Cebp-β (Figure 4a). S33, S37 and T41 can be modified by the enzyme GSK3beta[17–19], whereas S45 is modified by CK1[19]. Both inhibitor (Figure 4c) and BRD4 knock-down cell (Figure 4d) showed decreased levels of phospho-GSK. PDGF-BB induced increased phosphorylation of GSK, however, this effect was attenuated in BRD4 knock-down cells (Figure 4E, 4F). These suggest that tumorigenesis by PDGF-BB is mediated by BRD4, GSK, and β-catenin by attenuating β-catenin degradation in soft-tissue tumor.

**Figure 4.**
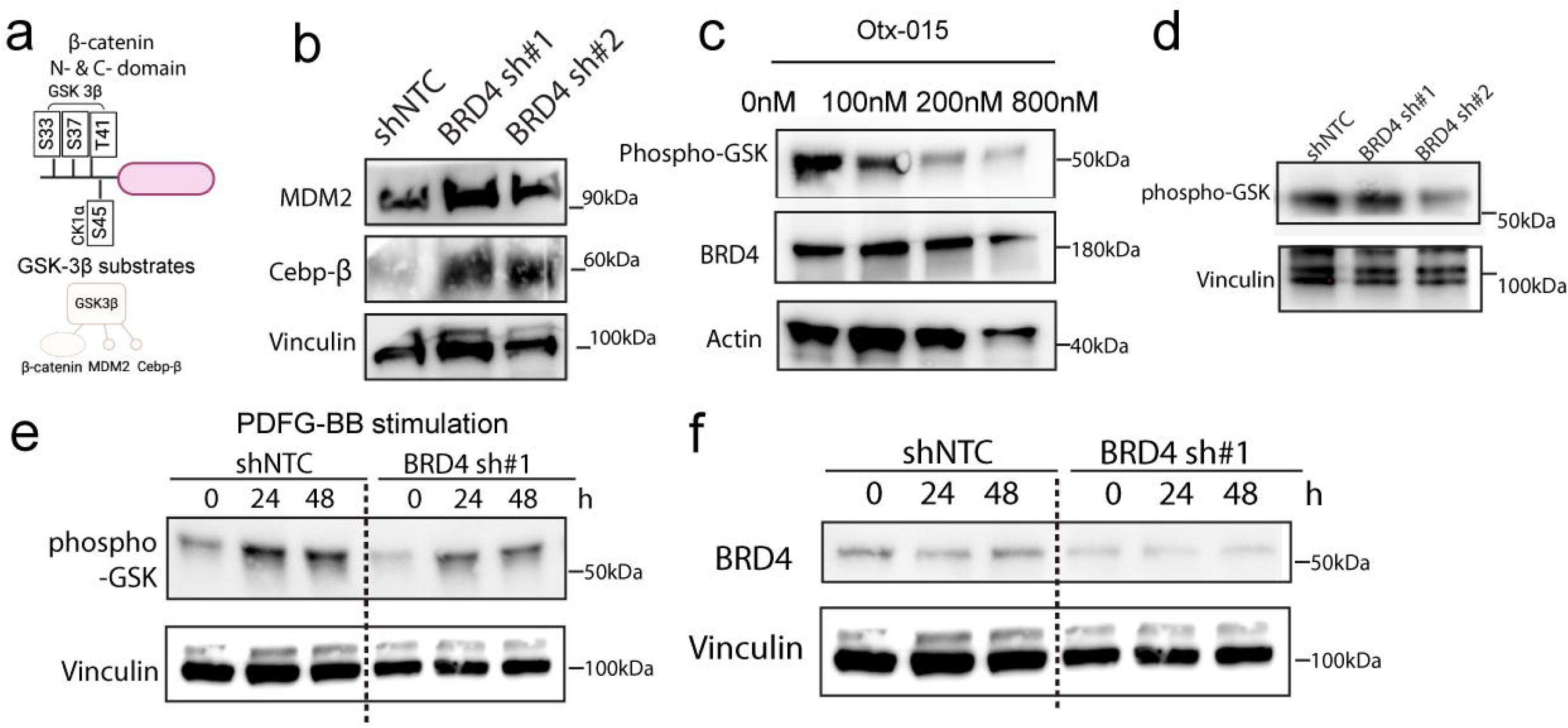
PDGF-BB requires BRD4 in regulating GSK to degrade β-catenin. β-catenin phosphorylation sites associated with degradation in its N-terminal. and GSK-3 β know substrates. (b). MDM2, and Cebp β expression. Blot represent as least two biological repeats. (c). Effect of inhibitor in inhibition of phospho-GSK at different dosage (0, 100, 400, 800nM). (d). phospho-GSK was decreased in BRD4 knock-down cells compared to control. (e) Phospho-GSK was increased in PDGF-BB stimulation (0,24,48 hours); Decreased phospho-GSK was observed in BRD4 knock-down cells. (f) BRD4 in control cells, and BRD4 knock-down cells with and without PDGF-BB in 0, 24, 48 hours treatment.

### (#5). PDGFR/GSK/β-catenin and active β-catenin signaling were mediated by BRD4

PDGF-BB stimulated PDGFR, TCF4, as well as the active β-catenin signaling(Figure 5a-d), however, these effects were attenuated by BRD4 knocked down. These suggest that PDGFR/GSK/ β -catenin and active β -catenin signaling are mediated by BRD4. Consistent with this, *BRD4*, *CTNNB1* gene expression were not responded to in BRD4 knock-down cells (Figure 5e). Inhibitor decreased the protein level of β-catenin, TCF4 (Figure 5f), confirming disruption of the functions of BRD4 attanuates β-catenin/TCF4 signaling. Overall, these data suggest that BRD4 also mediates PDGF-BB in regulating active β-catenin.

**Figure 5.**
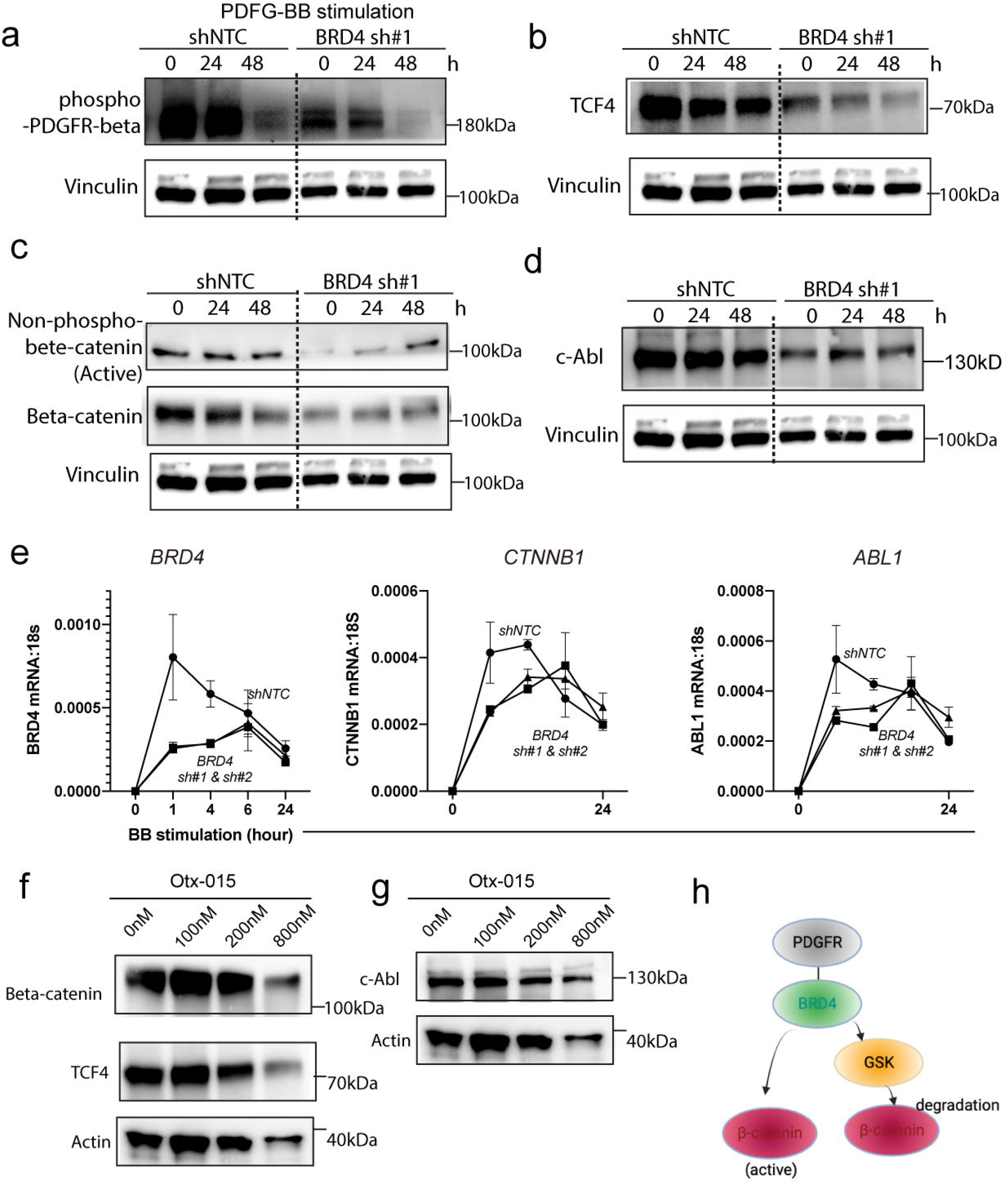
PDGF-BB regulated BRD4 mediated active β-catenin activation. (a-d) phospho-PDGFR, TCF4, Active beta-catenin, beta-catenin regulation in BRD4 knock-down cells, and control cells without and in the presence of PDGF-BB.(e)*BRD4*, *CTNNB1* and its target gene expression in the presence of PDGF-BB at 0, 4, 6, and 24-hour time points. (f-g)Gene regulated by inhibitors at different dosages (0, 100, 400, 800nM).

### (#6.) BRD4 regulated β-catenin through transcription from RNA polymerase II activity

To investigate how BRD4 regulated β-catenin, we perform the loss of function screening useing shRNA pool based on soft-tissue tumor cells, and cells treated with PDGF-BB (Figure 6a). As BRD4 was mainly maintained in nucleus, driver genes indentified in this screen with a cellular component GO terms associated with “nucleus” were presented (Figure 6b), and a string network with both BRD4 asssociated- and β-catenin associated - genes were ploted (Figure 6c). Figure 6d showed the details of these driver genes, and when BRD4 node-, and β-catenin node-genes were knocked down, how cell growth were manipulated (Figure 6e). Figure 6f-6i showed a subset of genes associated with both BRD4, and β-catenin were mapped with biological signature of “transciption from RNA polymerase II promoter” and “positive regulation of transcription (DNA-templated)”. Figure 6j summarized how BRD4 regulated both β-catenin degradation by GSK through transciption from RNA polymerase II promoter activity with PIK3C, and a potential way that BRD4 regulated nucleus active form of β-catenin through transciption from RNA polymerase II promoter activity, which contributes a non-canonical response from binding with β-catenin. Overall, these data strongly suggest that BRD4 regulates PDGF/β-catenin signaling through transcription from RNA polymerase II activity.

**Figure 6.**
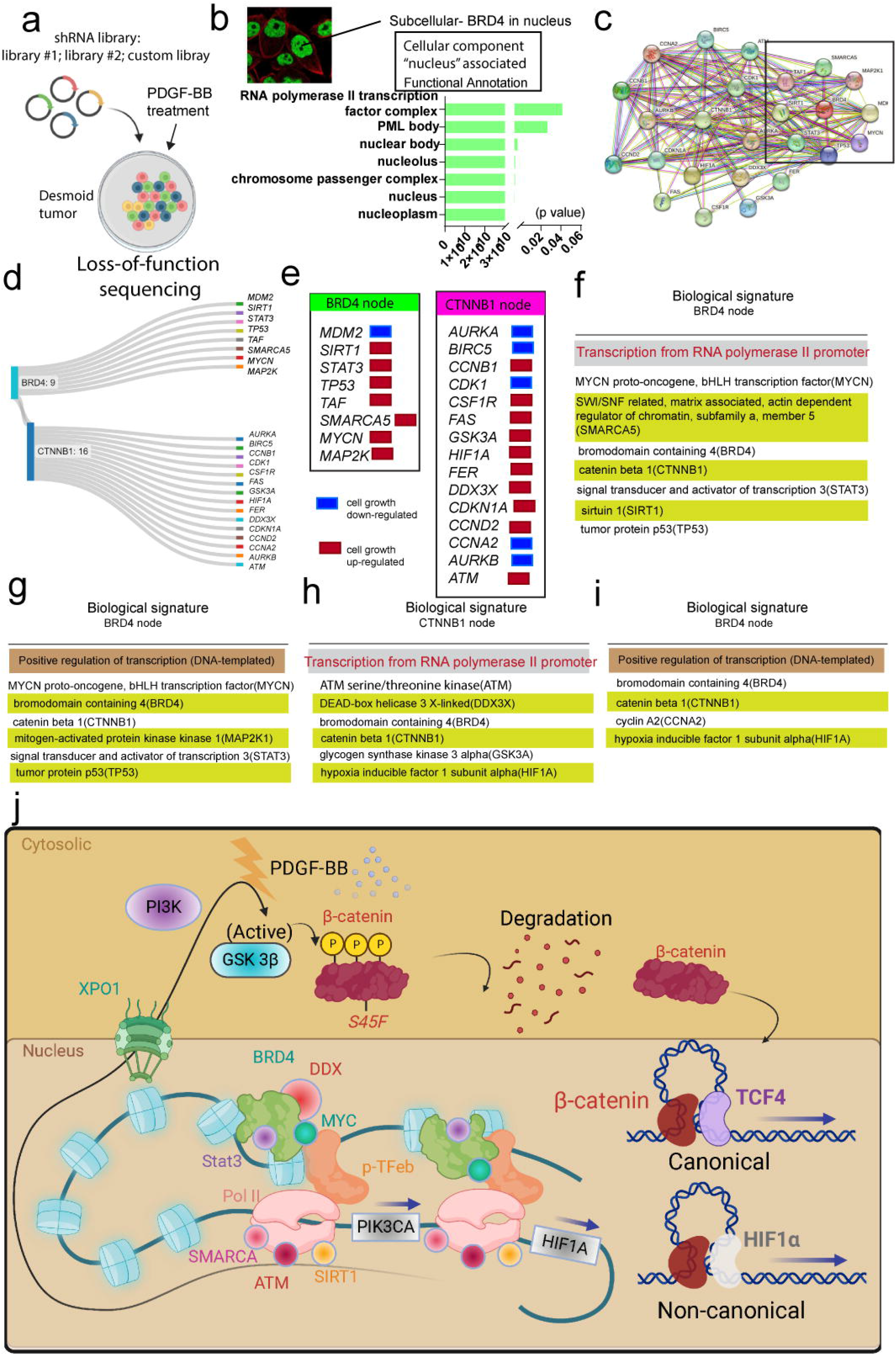
Loss-of-function screeen identifed BRD4 mediated PDGF-BB signaling in promoting β-catenin. (a)shRNA library and workflow with loss-of-function screening over soft-tissue tumor. (b) cellular component GO terms associated with “nucleus” over driver genes identified in the screen (c). String network with both BRD4 asssociated- and β-catenin associated - genes. (d, e) Details of driver genes associated with nucleus. (f-i). Genes associated with both BRD4, and β-catenin mapped with biological signature of “transciption from RNA polymerase II promoter” and “positive regulation of transcription (DNA-templated)”. (j)A proposed model to show how BRD4 regulated both β-catenin degradation by GSK through transciption from RNA polymerase II promoter activity with PIK3C, and a potential way that BRD4 regulated nucleus active form of β-catenin through transciption from RNA polymerase II promoter activity with HIF1a, which contributes a non-canonical response from binding with β-catenin, based on known references.

## Methods

### soft-tissue cell lines

Cell lines were grown according to the vendors’ instructions, were incubated at 37 °C in a humidified incubator with 5% CO2, and passaged every 3–4 days. For soft-tissue tumor BRD4 ko cells were stably infected with pLKP.1-BRD4 shRNA lentivirus.

### Isolating plasmid DNA for lentivirus production

*E. coli* cells were transformed with shRNA plasmid (hairpin-pLKO.1 vector with Puromycin resistance gene and the *MDM2, BRD4* shRNA). Packaging plasmid (pCMV-dR8.91) and envelope plasmid (pCMV-VSVG) and scramble control plasmid in the pLKO vector were prepared. Transformed bacterial cells on LB + Ampicillin Agar Plate were grown overnight at 37°C. Individual colonies were picked and grew 250ml of LB + Amp (100 ug/ml) broth and grew O/N (12-16 hrs.) at 37°C at 250 RPM. Bacteria was spun down to pellets in ultracentrifuge at 6000 x g for 15 min. at 4°C. Supernatant was removed. Pellet was put in −80°C for at least one hour to facilitate cell lysis. MaxiPrep with Qiagen HiSpeed Plasmid Maxi Kit was used. Plasmid DNA was quantitated.

### soft-tissue cell line mutation sequencing

Genomic DNA was isolated from cells. *CTNNB1* gene locus was amplified by PCR. 40 ng of DNA was added with primers per reaction, and then samples were put into the PCR machine. Agarose gel electrophoresis was used to confirm the product. The proper sized band was 529 bases for *CTNNB1* exon 3 amplicon. PCR products were purified and sent for Sanger Sequencing.

### Reverse transcription procedure for producing lentivirus in 293T cells

Hek293T cells were thawed in T175 and grew to ∼80-90% confluence (Hek293T media = DMEM HG + 0.1X P/S + 10% FBS). One T175 culture was split into eight 10 cm dishes. For each construct, we have two 10 cm dishes of Hek293T cells. Plasmid DNA-Lipofectamine 2000 (Invitrogen) complexes were prepared in a 10cm dish to dilute plasmids into 2mL of Opti Mem. Each dilution should contain pCMV-dR8.91, pCMV-VSVG, and the shRNA construct of interest (scramble, and knockout). Lipofectamine 2000 was diluted in Opti Mem in a separate 15 ml conical tube. Diluted DNA was combined with the diluted Lipofectamine 2000, and the combined plasmid DNA/Lipofectamine was incubated for 20 minutes at RT to allow DNA-Lipofectamine complexes to form. Hek293T cells were lifted off the 10 cm dishes and cells were resuspended in 6 ml of growth medium (DME HG 10% FBS without antibiotics) in a 15mL conical tube. 4ml of Lipofectamine-DNA complexes was added to a T75 tissue culture flask; 6 ml growth media was added into the flasks. After one day, the medium containing the Lipofectamine was removed, and replaced with 12 mL high BSA growth media/viral harvest media. Virus-containing supernatants were harvested for the first 48 hours by removing the harvest medium and were transferred into a 50 ml sterile conical tube. Cell debris was pelted and supernatant which contains the virus was taken by a cold centrifuge 1000 rpm x 5 min. Supernatant was filtered through 0.45 MIC Nalgene syringe filters using 30mL syringes into a new 50 mL tube. The virus was concentrated using Amicon Ultra filters (Millipore cat. no. UFC910024) until they have about 1.5ml of concentrated virus, and stored at −80°C.

### Infecting cell with virus

Cells should be 50-75% confluent. Polybrene was dissolved in PBS to make a Polybrene stock solution of 10 mg/ml. By creating a 16 ug/ml polybrene solution, the final concentration is 8ug/ml on the plate. FM media followed by Polybrene media was added to the cells, and then virus was added for experimental groups, and controls with no virus. Puromycin media was made with the desired final concentration of 1 µg/ml. Virus media was removed from plates, and 10ml of Puro media was added to each 10cm plate. Check after day 2 or day 3 after infection, and pick which clone has the best effect for our end goal.

### Cyquant Proliferation Assay

Cells were trypsinized and be prepared to count in either a 15mL or 50mL conical, depending on how many plates needed to harvest to do the experiment; 1000 cells per well were plated; Drug was added the next day, and Day 0 plate was collected. The 96-well plate was sealed, and was stored at −80C until the day to perform the Cyquant assay; All of the plates were collected at their indicated time points. The media was refreshed on day 3. On the day of assay, plates were removed from the freezer and let them thaw fully. Basic Cyquant Protocol was proceeded.

### Drug treatments of cells *in vitro*

For drug treatment of cells, 2.5 μM BRD4 inhibitors alone were added to triplicate wells to a volume of 100 ul. Control experiments were performed without the addition of drugs using the vehicle (DMSO) only. Day 0, day 2, day 4, and day 6 cells were collected to be measured for cell viability using the Cyquant method. Cell viability was also determined by trypan blue dye exclusion in some experiments. Apoptosis was assessed using Annexin V as per the manufacturer’s instruction.

### RNA extraction, cDNA synthesis, and qPCR

RNA was extracted using the RNeasy Mini kit according to the manufacturer’s protocol. On-column DNA digestion was carried out using DNase1. Eluted RNA was quantified using the Nanodrop 2000 and RNA was used in cDNA synthesis, according to the manufacturer’s instructions. qPCR was done using SYBR Taq master mix, and Probe of each primer of interest. The qPCR reactions were performed for melt curve analysis.

### Gene knockdown

Gene knockdown or overexpression in Ds cells was achieved by viral transfection. DNA constructs were used to generate a virus for infection of cells to generate knock-down or overexpression (pTRIPZ and pLKO.1 for knock-down). To generate the virus, 2 × 106 HEK-293T were plated one day before transfection at 2 million cells per plasmid in a 10 cm plate in culturing media. Transfection was carried out in antibiotic-free culturing media using lipofectamine in Opti-mem media. This was combined with a further Opti-mem containing plasmid DNA, ps PAX2, and VSVG, incubated at room temperature, and then added dropwise onto the target plate of HEK-293T. After the medium was aspirated and refreshed with normal culture medium containing antibiotics as above. Two days after transfection of the HEK-293T, infection of the target cells was initiated by spinning down target cells per sample and re-suspending in culture media. Virus-containing media was removed from the plate of 293 T cells, and mixed with polybrene. To each well of target cells, 1 ml of virus-containing media was added. A GFP-expressing virus and a ‘cells only’ well were included as controls for the transfection and infection respectively. Cells were then transferred to the flask with media for overnight incubation. The following day a second infection was carried out as described above, and after this, the cells were selected with puromycin.

### Statistical analysis

Results were statistically analyzed in GraphPad Prism (version 9.0, GraphPad Software Inc., San Diego, CA) and presented as mean±SEM. Statistically significant differences between the three, or two groups were assessed by a one-way ANOVA, and two-tailed unpaired t-test, respectively. p ≤ 0.05 was considered significant. ns, not significant; *p ≤ 0.05; **p ≤ 0.01; ***p ≤ 0.001; ****p ≤ 0.0001 for indicated comparisons. Statistical details of each experiment can be found in the Results and Figure Legend sections.

## Discussion

In this study, we uncovered the PDGF/GSK/β-catenin signaling as a crucial survival mechanism in soft-tissue tumor cells. PDGF-BB is a regulator of active β-catenin expression, and the activation of PDGF-BB promotes the interaction between β-catenin and TCF4, presumably through a BRD4/phospho-GSK dependent mechanism; Inhibition of BRD4 drives cell cycle arrest, and apoptotic response in tumor cells. These results support that targeting the PDGF-phospho-GSK-BRD4 pathway provides a novel strategy for treating soft-tissue tumors.

Activation of PDGFR signaling may be coupled with multiple downstream pathways in the regulation of cell growth, proliferation, migration and survival[15]. In tumor-associated endothelial and fibroblast stromal cells, PDGF has been shown to activate Akt- and MAPK-dependent survival mechanisms[21–24]. In this study, we provided molecular evidence demonstrating that in soft-tissue tumor cells PDGF could significantly promote nuclear translocation activity of β-catenin which is essential for cell survival. BRD4 depletion led to the inhibition of phospho-GSK expression, degradation of β-catenin, and induction of apoptosis. Our results are consistently with Iqbal et al. [25] that PDGF induce rapid phosphorylation of both Akt and glycogen synthase kinase 3-β (GSK-3β) in human prostate cancer, which may also contribute to the elevated intracellular levels and nuclear accumulation of β-catenin[26]. In human colon cancer cells, PDGF-BB induces EMT[27] and upregulates cyclin D1 and c-Myc[28] by activating β-catenin-dependent gene expression. PDGF-BB induces the phosphorylation of kinase, which subsequently recruits p68, an RNA helicase with ATPase activity, and activates its phosphorylation. Thus, there could be other pathways mediated by BRD4 targeting beta-catenin in PDGF-BB signaling.

Our data confirmed a highly active β-catenin/TCF4 signaling regulated by BRD4 in soft-tissue tumor cells, and BRD4 may regulate β-catenin target genes through TCF4. Constitutive activation of the Wnt/β-catenin pathway and upregulation of Wnt signaling target genes are common in soft-tissue tumors. The activating β-catenin mutations in soft-tissue tumor could stabilize β-catenin, resulting in nuclear β-catenin accumulation and the formation of the nuclear β-catenin/TCF4 complex [29]. The acetylation of β-catenin further promoted the association of β-catenin with TCF4, eventually activating the transcription of β-catenin target genes [29,30]. Although that human c-Abl promoter does not contain any consensus sequences of TCF/lymphoid enhancer-binding factor (LEF). These suggest that certain transcription factor(s), other than TCF, could be responsible for β-catenin activation of c-Abl transcription. Here, we found that the transcription factor was found to interact with β-catenin in soft-tissue tumor cells, which may mediate c-Abl transcription by binding to the promoter. These data are consistent with a previous study showing that β-catenin can switch its binding partner from TCF4 to HIF-1α and enhance HIF-1α-mediated transcription, and this dynamic reassembly of β-catenin with HIF-1α may allow colorectal cancer cells to rapidly adapt to hypoxic stress and survive[31]. PDGF may significantly affect the expression of β-catenin target genes identified by Chip-seq in our result by promoting the interaction between β-catenin and HIF-1α in a Wnt-independent mechanism. Further, our results also showed that BRD4 inhibition abrogated TCF4 function and down-regulated its direct target MYC. Therefore, this supports that BRD4, β-catenin and its downstream components should be important therapeutic targets for soft-tissue tumors. Regulation of bete-catenin activity by PDGFR-β and targeted therapies modulates soft-tissue tumor cell proliferation, suggesting a reason for variable biologic behavior between tumors and a mechanism for sorafenib activity in dermoid tumor.

The prevention of β-catenin degradation by BRD4, as a consequence, could provide pivotal protection against apoptotic signals in soft-tissue tumor cells. The p53 tumor suppressor is an important transcription factor that regulates cellular pathways such as DNA repair, cell cycle, apoptosis, angiogenesis, and senescence [32]. The increase in p53 levels triggers either cell cycle arrest or apoptosis[33]. Our results showed that BRD4 induces cell cycle arrest, apoptosis independently of p53 since BRD4 inhibition induced cell apoptosis stimulated less p53 proteins. It is suggested that GSK-3β phosphorylation at Ser9 inhibits its apoptotic activity, whereas phosphorylation at Tyr216 promotes its apoptotic activity[34]; Overall, this may suggest that BRD4 induces GSK-3β phosphorylation at Ser9 to inhibit apoptosis independent of p53 in soft-tissue tumor.

BRD4 as a prime therapeutic target that could potentially be effective in most soft-tissue tumors regardless of their underlying β-catenin genetic mutations. There are few studies associated between β-catenin and BRD4. Song *et. al* indicated that BRD4 expression is negatively correlated with the overall survival of gastric cancer patients, and knockdown of BRD4 attenuated the stemness of gastric cancer cells[35]. The role of BET1 inhibitors on metastatic capability was addressed by Jianlin Wang et al., and the prostate cancer cells with inhibitors acquired a spindle-shaped morphology with repressed invasion[36]. These results support our observation that BRD4 depletion has been shown to induce a more flattened and elongated shape of cancer cells, which is indicative of a more differentiated phenotype. It has been suggested that the effect may be related to alterations in the cytoskeleton and cell adhesion molecules, as well as changes in the expression of genes involved in cell differentiation and migration[35]. In absence of Wnt, β-catenin is usually degraded by destruction complex. Here, we showed that PDGF signaling promoted the degradation of β-catenin even though S45F is mutated, and other phosphorylation sites are important for degradation of β-catenin. This correlates with the fact that Wnt signaling primarily inhibits β-catenin phosphorylation at Ser33/Ser37/Thr41 by GSK3 but not Ser45 phosphorylation by CK1 [20, 37]; Overall, the observation that knock-down of BRD4 affects cell morphology in cancer cells highlights the potential of BRD4 inhibitors as a therapeutic strategy for most soft-tissue tumors regardless of their underlying β-catenin genetic mutations.

Collectively, our data revealed that aberrant BRD4 knockdown significantly decreases soft-tissue tumor aggressiveness and unfavorable prognosis in soft-tissue tumor when PDGF-BB was presented as a pro-oncogenic factor driving cell proliferation. PDGF-BB stimulates GSK through BRD4 with a regulation of transcription from RNA polymerase II activity, thereby initiating the β-catenin degradation, and BRD4 activated HIf1a in combination with nuclear activation of beta-catenin to promote cell proliferation. Our study delineated a novel signaling axis that may allow soft-tissue tumor cells to escape apoptosis during colonization by activating PDGF-GSK-β-catenin pathway in cancer cells. It is plausible to hypothesize that PDGF-BB may be crucial in mediating tumor and microenvironment. Specific targeting of both PDGF and BRD4 survival pathways in soft-tissue tumor cells could provide a new strategy to disrupt the vicious cycle and efficiently treat soft-tissue tumors.

## Ethics Statement

This article does not contain any studies with human participants or animals performed by any of the authors. For this type of study, formal consent is not required.

## Acknowledgement

Sincere gratitude to soft-tissue tumor cells which were immortalized by ectopic expression of *TERT* at Dr. A.M.C lab at MSK. The authors give thanks to contribution to the paper as follows: Technical help (methods: K. P; chemicals/reagents and resources: Dr. A. M. C).

